# Sex Differences in Structural and Receptor mRNA Expression in the Ventral Anterior Cingulate Cortex and a Potential Role of Perineuronal Nets in Monogamous Pair Bond Establishment (*Peromyscus californicus*)

**DOI:** 10.1101/2025.06.16.659986

**Authors:** Candice L. Malone, Jiaxuan Li, Elsa M. Luebke, Leykza Carreras-Simons, Warren W. Treis, Emma R. Hammond, Patrick K. Monari, Catherine A. Marler

## Abstract

The monogamous California mouse (*Peromyscus californicus*) exhibits distinct behavioral changes during pair bond formation. Using a detailed temporal behavioral analysis over seven days, we found a rapid decrease in aggression within 24 hours of pair introduction in this highly territorial species. After this aggression reduction, the gradual increase in affiliative behaviors varied by type of affiliative behavior and ranged from one to seven days. We then measured neurobiological changes at three time points during this transition to uncover mechanisms that might govern this shift from aggressive to affiliative behavior, revealing novel sex differences that add to current research on biological mechanisms of social bonding. Specifically, we examined plasticity through mRNA expression of two perineuronal net (PNN) associated proteins, HAPLN and ACAN, in two brain regions implicated in affiliation, aggression, and social cognition: the ventral anterior cingulate cortex (vACC) and lateral septum (LS). The vACC in females exhibited higher expression levels of both of these PNN components relative to males. Additionally, we observed a decrease in ACAN mRNA expression in the vACC over the course of pair bond establishment, but no such change in the LS. Furthermore, oxytocin receptor (OXTR) and vasopressin receptor (AVPR) plasticity exhibited sex-specific patterns in the vACC during pair bond formation. Females displayed higher OXTR mRNA expression across the bonding period, whereas males expressed higher AVPR mRNA levels. In addition, our results uncovered a positive association between AVPR expression levels in the vACC and male-female nesting distance, a novel measure of affiliative behavior. We discuss how a decrease in PNNs could allow for an increase in receptor plasticity in the vACC as the pair bond is established. Moreover, we suggest that structural plasticity across this social transition may differ between males and females due to factors such as pre-pair sociality and aggression/territoriality changes.

**Plain English Summary:** Some mammals, such as the monogamous California mouse, establish a long term partner with which they defend territories, forage, and raise young. This is a large behavioral shift, wherein these animals form a cooperative dyad. In order to behaviorally adjust to this social change, we expected to see neural changes that are indicative of this adjustment, such as changes in neurons that make them more flexible or responsive to social stimuli. While we detected an increase in neural plasticity across the pair bond for both sexes, we also noted important sex differences in cortical regions that were not expected. The anterior cingulate cortex, which is conserved across mammals, is a brain area associated with social decision making and empathy. Sex differences in both receptors and plasticity were detected in this area, indicating that the sexes may use different neural mechanisms to adjust to large social changes. Further exploration into how the sexes use different modes of neural plasticity to appropriately adjust behavior is needed to explain this novel finding.

**Highlights:**

1. California mice exhibited a graded shift from male-female aggression to greater affiliation across pair bond establishment.
2. Aggrecan mRNA expression, an indicator of perineuronal net (PNN) abundance, decreased in the ventral anterior cingulate cortex (vACC), but not the lateral septum (LS), across pair bond establishment.
3. Females displayed greater aggrecan mRNA in the vACC relative to males.
4. The vACC displayed greater oxytocin receptor (OXTR) mRNA expression in females and greater vasopressin receptor (AVPR) mRNA expression in males.

## 1.1 Introduction

The monogamous California mouse exhibits a clear behavioral change during pair bond formation (Ribble, 1991). Both males and females exhibit high levels of hormonally-modulated territorial aggression, which is dampened during the process of initial pair bonding (Kuske et al., 2024; Kleiman, 1977). Behavioral transitions, which can be defined by bouts of social learning and behavioral change, are important for appropriate context-dependent behaviors. The mechanisms underlying these behavioral transitions are expected to be reflected in changes in the restructuring of receptor densities (Ophir et al., 2012) and even in changes of extracellular matrix proteins of neurons (Cornez et al., 2018). California mice are thought to display particularly selective partner selection (Gleason et al., 2012). This selectivity is exemplified in the long establishment period, which may last at least seven days in this species, indicated by an increase in the duration of affiliative vocalizations across this time (Pultorak et al., 2018; Pultorak et al., 2015). This long establishment period may contrast with the monogamous prairie vole which can mate and bond within 24 hours (Meek et al., 1993). California mice, on the other hand, appear to have longer latencies to mating and higher variation in mating time, and do not bond within 24 hours (Gleason et al., 2010; 2012; Kowalczyk et al., 2018). This is particularly true in larger housing arenas in which birth latencies can range from approximately 30 to 90 days (Gleason et al., 2010), but matings can also occur within 24 hours when the paired female is in estrus or in small, less naturalistic housing cages (Roy et al., 2025). The biological mechanisms that govern an observable transition from aggressive to affiliative behavior have not been readily studied. In the current study, we assessed both neuropeptide plasticity in a cortical (vACC) and subcortical brain region (LS) and the time course for changes in neuropeptides across the relatively slow and multiple behavioral changes during pair bond formation in California mice. Additionally, we explored changes in measures of plasticity through measures of perineural nets as a measure of structural plasticity.

Receptor plasticity has been repeatedly studied in the investigation of pair bonding and monogamy in prairie voles (Hirota et al., 2020; Aragona et al., 2004, for review see Young & Wang, 2004). Two structurally similar neuropeptides, oxytocin (OXT) (Fricker et al., 2023) and vasopressin (AVP) (Tomaszycki et al., 2017), have been compared between unpaired and paired monogamous prairie voles. In paired individuals, these neuropeptides and their receptors tend to be more highly expressed. OXT and AVP have been implicated in both aggressive and affiliative behaviors across animal taxa (Donaldson et al., 2010; Klatt et al., 2013; Jiang et al., 2018). OXT receptor density increases in reward-related brain regions such as the nucleus accumbens and caudate putamen during pair bonding in both male and female prairie voles (Ophir et al., 2012; Young & Wang, 2004). However, the dynamic temporal changes of OXT receptor density over pair bond formation have not been examined.

AVPR, in contrast, has long been associated with aggressive displays in rodents both via mating-induced aggression and territoriality. Central and peripheral AVP administration facilitates the display of territorial aggression in adult male hamsters (Ferris & Potegal, 1988), while the blocking of these receptors in the anterior hypothalamus reduces aggression (Ferris et al., 1990). More AVP+ cells in the anterior hypothalamus are associated with higher numbers of aggressive displays in males (Ferris et al., 1989). Sex differences in the expression of AVPR are also present throughout subcortical areas including the hypothalamus (Dubois-Dauphin et al., 1996). In the California mouse, peripheral and central AVP administration increases aggression (e.g. wrestling) in males and females (Guoynes & Marler, 2021, unpublished; Bester-Meredith et al., 2005). Despite the sex specific functions of these neuropeptides, both OXT and AVP are associated with either affiliation or aggression depending on the social context (Oliveira et al., 2022; Bosch et al., 2010; Bosch et al., 2012; Schmidt et al., 2018; for review see Nephew et al., 2013; Neumann et al., 2012). OXT is also thought to be more influential on female social behavior, whereas AVP is thought to be more influential on male social behavior (Dumais & Veneema, 2016). This sex-based difference in utilization of OXT and AVP persists in studies of pair bonding behavior (Hammock et al., 2006). The context-specific nature of OXT and AVP effects may be modulated by changes in the proportions of OXT to AVP receptors in various brain regions involved in sociality. Interestingly, OXT, especially in the dorsal LS, is noted to modulate anxiety-related behaviors, which may mediate the context-specific social effects of its administration (Pirri et al., 2025; Grillon et al., 2013).

The subcortical region we explored, the LS, is one component of the social neural network (O’Connell & Hoffman, 2011) and is highly implicated in the gating control of aggression and affiliation, especially in female rodents (de Moura Olivieria et al., 2021). The ratio of AVP to OXT expression in the LS can alter the social behavior displays of rats (de Moura Olivieria et al., 2021). Chemogenetic activation of the LS increases affiliative bonding behavior, social approach, and extra-pair aggression (Sailer et al., 2022). The LS acts as an integration center for multiple modes of environmental and social context cues, which allows for greater accuracy in appropriate behavioral displays (Menon et al., 2022). Furthermore, the LS can be divided into multiple subdivisions, with sex-specific differences in steroid and peptide receptor expression, connectivity, and size, already noted in the dorsal LS and ventral LS (Tsukahara et al., 2002; 2004; 2014; De Vries et al., 1981; 1983;). This demonstrates that the LS plays an important role in navigating socially appropriate displays of both affiliation and aggression (Menon et al., 2022). OXT and AVP receptors within the LS may be central to this balance.

Connected to the LS, the ventral anterior cingulate cortex (vACC) is an area in the medial prefrontal cortex repeatedly implicated in the control of social behaviors across species, including humans (reviews by Heukelum et al., 2021; Li et al., 2021). The vACC has known involvement in prosocial and aggressive behaviors (Basile et al., 2020; Heukelum et al., 2021; Li et al., 2021; López-Gutiérrez et al., 2021). In addition, the ACC has repeatedly been investigated in rodent models of empathy (review by Kim et al., 2019), and social decision making tasks (Zhong et al., 2017; Apps et al., 2016). ACC to LS connectivity has been implicated in the control of context-specific grouping strategies and social investigation (Fricker et al., 2024). OXT and AVP in the ACC in particular have been implicated in complex social behaviors, such as helping a familiar conspecific escape a negative stimulus or consoling familiar conspecifics after stressful events (Yamagishi et al., 2020; Huang et al., 2024). However, the complexities of OXT and AVP action in the vACC are far less studied than the LS.

While receptor plasticity is likely to change as a function of social experience (e.g. pair bonding), a more structural form of plasticity may also be influenced by social experience, such as perineuronal net (PNN) fluctuations in the cortex. PNNs are concentrations of protein scaffolds that gather around specific neurons after they have undergone synaptic or dendritic changes (Köppe et al., 1997; Celio et al., 1998; Cornez et al., 2018; Wang & Fawcett, 2012; Karetko & Shaniel-Kramska, 2009; Bosiacki et al., 2019). As such, they are thought to solidify these changes, inducing long-lasting learning and behavioral alterations. PNNs typically form around parvalbumin-positive neurons, and the presence of parvalbumin neurons can indicate the start of a transition period, whereas PNN formation can indicate the end of such a transition (Takesian & Hensch, 2013). Across the songbird literature, the formation of correct birdsong is often studied as a model of social learning (Nelson et al., 1994; Nottebaum et al., 1984). Song crystallization corresponds to increased PNN numbers in the song nuclei of songbirds (Cornez et al., 2018), indicating that PNNs may play a role in facilitating appropriate adult song production and learning. Furthermore, PNNs are found in the LS in rodents and other mammals (Seeger et al., 1994; Brauer et al., 1993; Brauer et al., 1995; Brückner et al., 1996). PNN formation has also been noted in other cortical areas, such as the ACC, which is associated with the control of social behavior (Heukelum et al., 2021; Yamagishi et al., 2020; Alpár et al., 2006). Recent literature in the California mouse has evaluated PNN changes in the medial prefrontal cortex, medial amygdala, and the medial preoptic area as a result of social exposure to pups (Acosta, 2024). As such, we evaluated the PNNs in the vACC and LS using immunohistochemistry and qPCR. We proposed that behavioral changes associated with the formation of long-term pair bonds in monogamous species are also supported by these PNNs.

We conducted a well-documented, longitudinal behavioral characterization of the early pair bond (similar to Roy et al., 2025) with three time points of tissue sampling to link with the receptor and neural plasticity changes. We hypothesized that OXT and AVP receptor densities in cortical (vACC) and subcortical areas (LS) would change across pair bond establishment, although we had no a priori predictions regarding the directionality of this change. We predicted, however, that females would display higher expression of OXTR and males higher levels of AVPR in both brain areas, and greater sex differences would be expressed in the LS, as this area is sexually dimorphic (Menon et al. 2022; Liu et al., 2012; Markham et al., 2002).

Alongside this prediction in receptor plasticity alterations, we predicted that decreases in PNN density and number would occur across the pair bonding period as measured by both qPCR and immunohistochemistry. Both the vACC and the LS not only contain PNNs, but are also overlapping nodes in both aggressive and affiliative behavior and contain OXT and AVP receptors (Ko, 2017), although interactions between PNNs and neuropeptide receptors remain unclear. We also predicted that vACC-PNN and LS-PNN formation, as quantified by cell number via IHC and visualization using confocal microscopy, would be more prominent in unpaired animals versus paired due to PNN’s tendency to decrease during critical periods of social learning (Cornez et al., 2018). Given that forming long-lasting pair bonding is a time of great change and behavioral adjustment, we predicted that after forming a full pair bond, both males and females of a pair would display greater numbers of PNNs, indicating such a learning period. We predicted no sex differences in vACC-PNN, but acknowledged that due to the differing roles of the LS in males and females (Oliveria et al., 2019; 2021) a sex difference might occur. However, we had no directional hypotheses regarding these sex differences.

## 1.2 Methods

We worked with 33 pairs of bonded male and female adult California mice (*Peromyscus californicus*) derived from the breeding colony at UW-Madison. Animals ranging from 3-9 months were used to form age-matched pairs. Animals were housed in an enriched environment and in large, acrylic cages (93 x 45 x 45cm) with open-air screen lids for the duration of the experiment. Mouse pairs were housed with two small, plastic animal igloo hideouts for nesting (10 × 10 x 8 cm), aspen bedding, two cotton balls for nesting, and food and water ad libitum. Mice were maintained at a 14:10 light:dark cycle, with all experimental observations taking place within 5 hours of the start of the dark phase. Sibling pairs were not used. Animals were maintained according to the National Institute of Health Guide for the care and use of laboratory animals. Procedures used in this study were approved by the University of Wisconsin– Madison College of Letters and Sciences Institutional Animal Care and Use Committee (Protocol L005447).

### 1.2.1 Behavioral Observations

After the start of the dark phase, animals were paired (n=66) on Day 0 of experimentation and observed and recorded for a 10-min period, which included initial introduction. Prior to pairing, all female mice were tail marked for visual identification. On successive days (Days 1-7), animals were video/audio recorded for 10-minute periods in the first 5 hours of the dark cycle. Prior to this recording, pairs remained in their home cage but moved to a recording room and allowed to acclimate for 10 minutes, as in previous lab protocols (Pultorak et al., 2018). Pairs were then separated for 5 minutes and reintroduced in their home cage to elicit vocalizations (Guoynes & Marler, 2021, unpublished observations). Prior to behavioral recording and immediately after the start of the dark cycle, nesting distances were evaluated with a tape measure and distance was recorded between the center of each nest. All other behavioral measures were coded using video recordings and the behavioral coding software BORIS (Friard et al., 2016). Nesting distance was evaluated each day prior to behavioral testing. Ultrasonic vocalizations (USVs) were recorded during behavioral testing using an ultrasonic microphone (Knowles FG).

An affiliative index was created from social grooming and huddling across the pair bonding period. These behaviors were normalized (z-scored) and summed to create the index. Behavioral indexes of behavior have previously been used to better grasp complex behavioral phenotypes (Malone et al., 2023).

### 1.2.2 USV Recording and Analysis

USV recordings of the 10-minute behavioral observations were analyzed using California Mouse Vocalization Neural Networks curated from our laboratory’s USV recordings over 10 years. These networks were trained using DeepSqueak (Coffey et al., 2019; version 3; available at github.com/DrCoffey/DeepSqueak), which was created to analyze rat and house mouse vocalizations. These networks can detect and categorize the four discovered vocalizations associated with the California Mouse: Complex sweeps, simple sweeps, barks, and sustained vocalizations (SVs) (Kalcounis-Rueppell et al., 2018; Pultorak et al., 2018; Pultorak et al., 2015). Training and model creation of these neural networks used over 20,000 sweep vocalizations and hundreds of SVs and bark vocalizations gathered from past experiments (Pultorak et al., 2018; Pultorak et al., 2015; Rieger & Marler, 2018; Rieger et al., 2021; Monari, Hu, Karnati, Malone, Xue, Zhu, Hammond, Li, Jang, Chen, Carreras-Simons, & Marler, unpublished). As noted in previous literature, these vocalizations have been associated with different social behaviors (Pultorak et al., 2018; Pultorak et al., 2015; Rieger & Marler, 2018; Rieger et al., 2021; Monari, Hu, Karnati, Malone, Xue, Zhu, Hammond, Li, Jang, Chen, Carreras-Simons, & Marler, unpublished). Specifically, barks and short SVs have been associated with aggressive encounters, while sweeps and long SVs are associated with higher levels of affiliative behaviors (Briggs & Kalcounis-Rueppell, 2011; Pultorak et al., 2015; Rieger & Marler, 2018). The current investigation differed from previous studies because it focused on the second harmonic of sustained vocalizations in order to reduce background noise below 20kHz, but may also have decreased ability to detect variation in SVs as previously found (Pultorak et al., 2015; 2017; 2018). This background noise is often produced by the mice when moving though aspen bedding. To investigate if sweeps and SVs differed in frequency, duration, or count across the pair bond establishment period, we analyzed USV recordings from Days 0, 1, 3, and 7. We evaluated the number, duration, and average frequency of each call type across days.

### 1.2.3 Experimental Procedure

For the qPCR study, subjects (pairs) were randomly assigned to one of three groups based on day after pair bonding: **Group 1** (n=10): behaviorally tested on day 0 and 1 and euthanized for brain extraction on day 1, **Group 2** (n = 10): behaviorally tested on day 0-3 and euthanized for brain extraction on day 3, or **Group 3** (n = 13): behaviorally tested on days 0-7 and euthanized for brain extraction on day 7. For the IHC investigation, an additional six sex-balanced unpaired controls, which were used as a comparison group for IHC, were housed identically to paired individuals, but were never exposed to a same-sex prospective pair-mate and were comparable to those sacrificed on day 1.

### 1.2.4 Real-time quantitative polymerase reaction (RT-qPCR)

Two biological assays, real-time quantitative polymerase chain reaction (RT-qPCR) and immunohistochemistry (IHC), were conducted with the tissue extracted after the longitudinal behavioral observations. We conducted both investigations for two reasons (1) We were interested in the subdivisions of LS but were unable to dissect subdivisions and evaluate them via qPCR and (2) Using IHC allowed us to confirm the localization of PNN+ cells in California mice, which has not been previously characterized, but has been investigated in other species (e.g. Cornez et al., 2018, 2020).

52 brains were used for RT-qPCR, to measure mRNA expression of OXT receptor (OXTR), AVP receptor 1a (AVPR), hyaluronan and proteoglycan link protein 1 (HAPLN), and aggrecan (ACAN). Areas of the vACC and LS were dissected using Fine Science Tools Sample Corer (Item No. 18035-02; Foster 251 City, CA, USA), placed in a 1.5ml microcentrifuge tube, and stored at -80C. Neural tissue was homogenized, and RNA was translated into cDNA prior to RT-qPCR. ACAN, HAPLN, AVPR and OXTR mRNA concentrations were evaluated via four primer pairs that were validated in the Marler lab (for more information, see supplemental note) and normalized to a validated housekeeper Rpl13a (Table 1) (Cuarenta et al., 2020). All samples were run in triplicate on a C1000 Touch Thermal Cycler (CFX96 Real-Time System, Bio-Rad, Hercules, CA) with PowerUp SYBR Green Master Mix (Thermo Fisher Scientific). Standard curves were used to evaluate efficiencies, which were between 95-105%. A single melt peak was used to determine the specificity of primer sets. The cycling steps were: denaturation at 95 °C for 2 min, followed by 44 cycles of 95 °C for 15 seconds (denaturation) and 58 °C for 30 seconds (annealing). Primer sets were developed using the NCBI Databases with associated accession numbers listed in Table 1. For more information on primer design, see supplemental. qPCR output was analyzed via ddCT method and represented as gene fold expression (2^-ddct). ddCT values were normalized to male and female “controls” or unpaired individuals. Rpl13a was used as a housekeeper gene for this experiment due to its stability across sexes and steroid hormone changes (Shah et al., 2011).

**Table 1.**
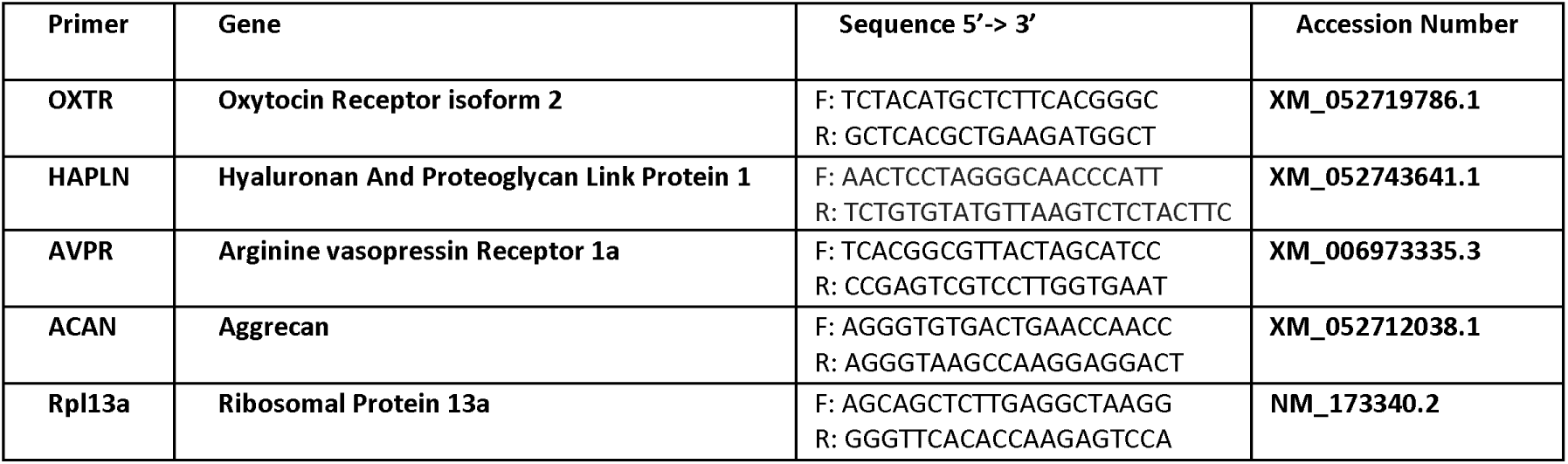
Details on primer sequence, gene target, and reference genome accession number.

### 1.2.5 Immunohistochemistry (IHC)

IHC was performed with 12 brains to verify general patterns of the number of PNNs in the ACC and LS in paired and unpaired control animals. Using a double-labeling immunohistochemical protocol on 40 micron slices (1-2 slices per brain), Parvalbumin positive (PV) and Wisteria floribunda lectin (WFA) antibodies were used to indicate PNN concentrations, as previously described by Cornez et al., (2018). We used a biotinylated primary antibody for WFA+ (Vector Labs; Newark, CA) alongside a secondary amplifying antibody for visualization (Alexa Fluor Anti-Streptavidin 488; Thermofisher), and a biotinylated primary antibody to label PV+ neurons (Successful IHC yielded identification of PNNs (WFA+) around LS PV+ (Parvalbumin Polyclonal Antibody; Thermofisher) with an amplifying secondary antibody for visualizing (Alexa Fluor Anti-Streptavidin 568; Thermofisher) cells in California mouse tissue, alongside a Hoechst nuclear DNA stain (Hoeschst 33342; Invitrogen, Fisher Scientific). We imaged brain slices with a Zeiss LSM 710 confocal scope using a 20x objective. Imagej software was used to process images. Each image was divided along a 100 micron grid, and five random grid squares were used for total cell counts.

### 1.2.6 Statistical Analyses

To evaluate the change in affiliative and aggressive behavior across pair bond establishment, we conducted a series of linear models and linear mixed effects models (repeated measures) using the General Linear Model (GLM) framework. We evaluated the effect of day as a fixed effect on the outcome variable for behavior (e.g. huddling, contact time, social grooming duration, wrestling duration, chasing, and olfactory investigation, nesting distance) with a random effect of pair identity. Dunnett’s multiple comparison corrected pairwise tests were conducted between behavior on Day 0 compared to all other days. An affiliative index was created from social grooming and huddling across the pair bonding period. These behaviors were normalized (z-scored) and summed to create the index. Linear models were used to evaluate the change in sweeps and SVs across the seven days, and Tukey’s multiple comparison tests were used to evaluate pairwise comparisons. Effect sizes were calculated using the effect size package in R/Rstudio. Effect size cutoffs are listed in Supplemental Material.

Two-way ANOVAs were used to evaluate the effect of pair bonding day and sex on relative gene expression. Due to violations of heteroscedasticity and non-normal residual spread, all gene expression values were transformed with a Box Cox transformation (lambda=0.5). This transformation resolved assumption violations. The OXTR/AVPR Ratio in the LS was not transformed, as it did not violate assumptions for a reliable two-way ANOVA test. Visualizations below use the non transformed gene expression values for easier interpretation, but the significant effects are derived from transformed data. Pearson correlation coefficients were conducted on non-transformed data.

To evaluate the correlative relationships between social behavior and mRNA expression, we created four correlation matrices divided by pair bond day and sex. We adjusted the significant p-value cutoff for each matrix using a Benjamini Hochberg correction (Menyhart et al., 2021). We also adjusted the p-value cutoff for significant correlations between behaviors and vocalizations (see Supplemental Fig. 1). For the relationship between vACC OXTR/AVPR and the affiliative index of behavior (Fig. 4D), which was significant after the cutoff adjustment, we noted an extreme outlier detected via Grubbs’ Method (alpha= 0.0001). After removal, the effect was no longer significant and was disregarded as a nonsignificant correlation.

Welch’s unpaired t-tests, which control for differences in variation between groups, were run between unpaired and paired animals in our IHC group. Welch’s unpaired t-tests were conducted between males vs. females when comparing PNN+ cells between sexes.

## 1.3 Results

### 1.3.1 Shifts from Aggressive to Affiliative Behavior

#### Aggression Rapidly Decreases After Pair Introduction

Chasing behavior was significantly different as a function of pair bond day (F (1.315, 21.61) = 3.946, p = 0.050, partial η2 = 0.19), with the incidence of this behavior decreasing over time. An outlier analysis was conducted due to the high variability in chasing behavior. This model was completed after removing 4 significant outlier observations from 106 observations from different subjects using Iterative Grubbs method with an alpha cutoff of 0.0001 (Rosner, 1983). We detected significantly more chasing behavior on Day 0 than on Day 2 (adj. p = 0.041) and Day 3 (adj. p = 0.040). Wrestling behavior did not display a significant effect as a function of pair bond day (F (1.910, 33.84) = 0.8160, p = 0.446). Chasing and wrestling behavior were positively correlated (R=0.34, p <0.001; Supplemental Figure 1).

#### Affiliation Gradually Increases Across Bond Establishment

Social grooming significantly increased across the pair bond establishment period (F (4.131, 73.17) = 3.199, p = 0.017,partial η2 = 0.15) (Fig. 1). A significant corrected pairwise comparison between Day 0 and Day 3 was detected (adj. p = 0.028). In addition to social grooming differences, a significant increase in huddling behavior across pair bond days was observed (F (4.247, 75.22) = 6.067, p < 0.002, partial η2 = 0.26). Significant pairwise comparisons were seen between Day 0 and Day 1 (adj. p = 0.006), Day 0 and Day 2 (adj. p = 0.003), Day 0 and Day 3 (adj. p = 0.001), Day 0 and Day 5 (adj p = 0.027) and Day 0 and Day 6 (adj. p = 0.044). No other significant pairwise comparisons were detected. Both huddling behavior and social grooming displayed an inverted U distribution, as noted by the nonsignificant pairwise comparisons between Day 0 and Day 7. Both behaviors tended to peak at day 3 of experimentation. The affiliative index (Fig. 1C) was positively associated with its components (social grooming and huddling duration) (R =0.90, p<0.001; R=0.90, p<0.001) and negatively correlated with contact bouts (R=0.36, p<0.001). Nesting distance significantly decreased as a function of day of pairing (F (1.893, 29.02) = 12.61, p <.001. partial η2 = 0.45) (Figure 1D).

**Figure 1.**
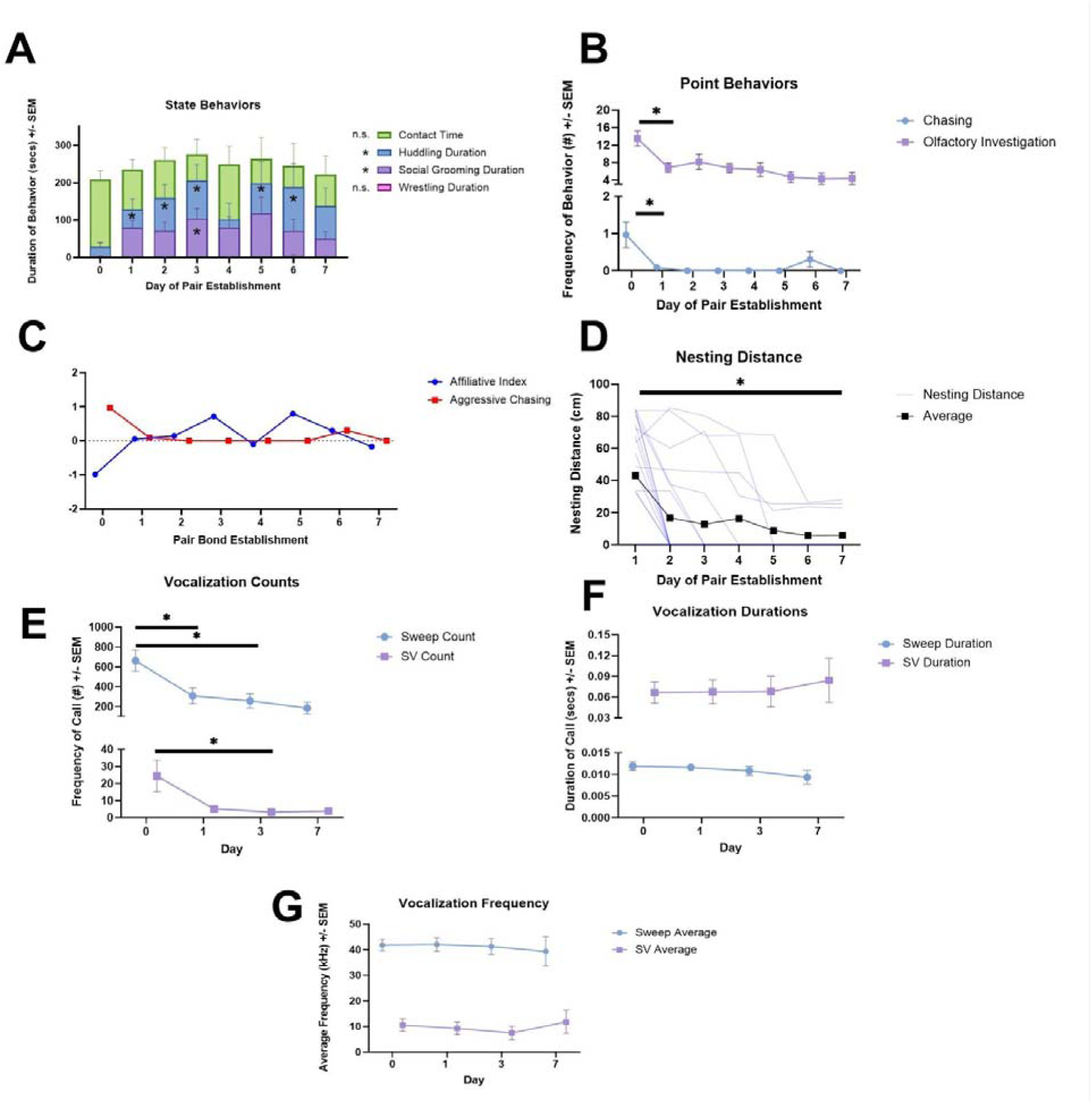
Across the pair establishment period, pairs experienced rapid reductions in aggression, followed by variable and gradual increases in affiliation. **A)** Both affiliative and aggressive state behaviors depicted with stars indicating significant pairwise tests when compared to Day 0. Both huddling and social grooming displayed an inverted U distribution across pair establishment days with significant increases but with no difference between Day 0 and Day 7. **B)** Both aggressive and investigative point behaviors decreased after pair introduction, with significance stars denoting the significant drop in both chasing behavior and olfactory investigation after Day 0. **C)** A clear decrease in agonistic tendencies is displayed (red line) from Day 0 to Day 1. An affiliative index (z-scored (huddling) + z-scored(social grooming)) showed gradual increases from Day 0 to Day 3. **D)** Individual pair’s nesting distances (cm) (blue lines), significantly decreased in a linear pattern across the seven days. **E)** Significant reductions in the number of sweeps and SV frequency were detected by Day 3. **F)** No significant changes in vocalization durations were detected across pair bond establishment, and **G)** no significant changes occurred in average vocalization frequency. * = P < 0.05.

#### Other Social Behaviors

Olfactory investigation significantly decreased as pair bonding progressed (F (4.129, 73.14) = 6.456, p < 0.001, partial η2 = 0.27). Dunnet’s multiple comparison corrected pairwise comparisons revealed that Day 0 olfactory investigation duration was significantly higher than all other days of evaluation (Comparison to Day 1: adj p < 0.003, Day 2: adj p = 0.032, Day 3: adj p = 0.0032, Day 4: adj p = 0.018, Day 5: adj p = 0.004, Day 6: adj p = 0.003, and Day 7: adj p = 0.016). Contact time duration did not differ significantly across pair bond establishment (F (3.970, 70.32) = 1.091, p = 0.367).

#### Vocalizations Decrease Over Pair Bond Establishment

The frequency of both SVs and sweeps decreased over time after pair introduction (Day 0), respectively (F(1.836, 37.95) = 6.275, p = 0.005, partial η2 = 0.23; F(1.068, 22.07) = 3.163, p = 0.087, partial η2 = 0.13)). Tukey’s multiple comparison tests of sweep counts across days revealed a significant reduction in sweeps between Day 0 vs. Day 1 (adj. p = 0.0365) and Day 0 vs. Day 3 (adj p = 0.014; Fig. 1E). For SV counts across days there was also a significant reduction between Day 0 vs. Day 3 (adj. p = 0.042; Fig. 1E). Linear mixed effects models were also used to compare changes in the average frequency and duration of both sweeps and SVs across pair bond establishment, but no significant differences were detected (Fig. 1F-G).

After performing a Benjamini Hochberg p-value correction, sweep counts were significantly and positively associated with SV counts, higher SV frequencies, longer SV durations (R = 0.50, p < 0.001; R = 0.50, p < 0.001; R = 0.38, p < 0.001). The average duration and frequency of sweeps were also positively linked (R = 0.56, p < 0.001). Sweep counts were also positively associated with contact bout number and olfactory investigation duration (R = 0.34, p < 0.001; R= 0.57, p < 0.001). SV counts were significantly and positively associated with higher average SV frequencies and longer SV durations (R = 0.47, p < 0.001; R = 0.45, p < 0.001). SV counts were significantly associated with chasing bouts and olfactory investigation (R = 0.63, p< 0.001; R= 0.44, p < 0.001). The average frequency of SVs was also positively associated with wrestling duration (R = 0.28, p = 0.005). In summary, across the pair bond establishment period, vocalization features were strongly correlated with social investigation behaviors (e.g. contact bouts and olfactory investigation). Namely, both sweep and SV counts appeared more commonly in bouts of heightened social investigation (see Supplemental Fig. 1). This aligns with previous literature in which sniff number was positively associated with sweep number (Pultorak et al., 2018). No significant correlations were detected between vocal features and biological measures from the vACC or LS.

### 1.3.2 qPCR Results

#### vACC-HAPLN and vACC-ACAN

Females, on average, displayed higher HAPLN mRNA expression levels in the vACC when compared to males in a nonsignificant trend with a medium effect size (F (1, 34) = 3.130, p = 0.086; partial η2 = 0.08 Fig. 2B). However, no other main effects or interactions were significant in this analysis of HAPLN (see Supplemental Table 1 for nonsignificant effects). For vACC ACAN mRNA expression, two significant main effects were detected: both a main effect of pair bonding day (F (2, 32) = 3.926, p=0.030, partial η2 = 0.20) and a main effect of sex (F (1, 32) = 9.799, p = 0.004, partial η2 = 0.23 Fig. 2A) were detected. Females, on average, displayed higher expression levels of ACAN and HAPLN (nonsignificant trend) across the entire pair bond establishment period.

**Figure 2:**
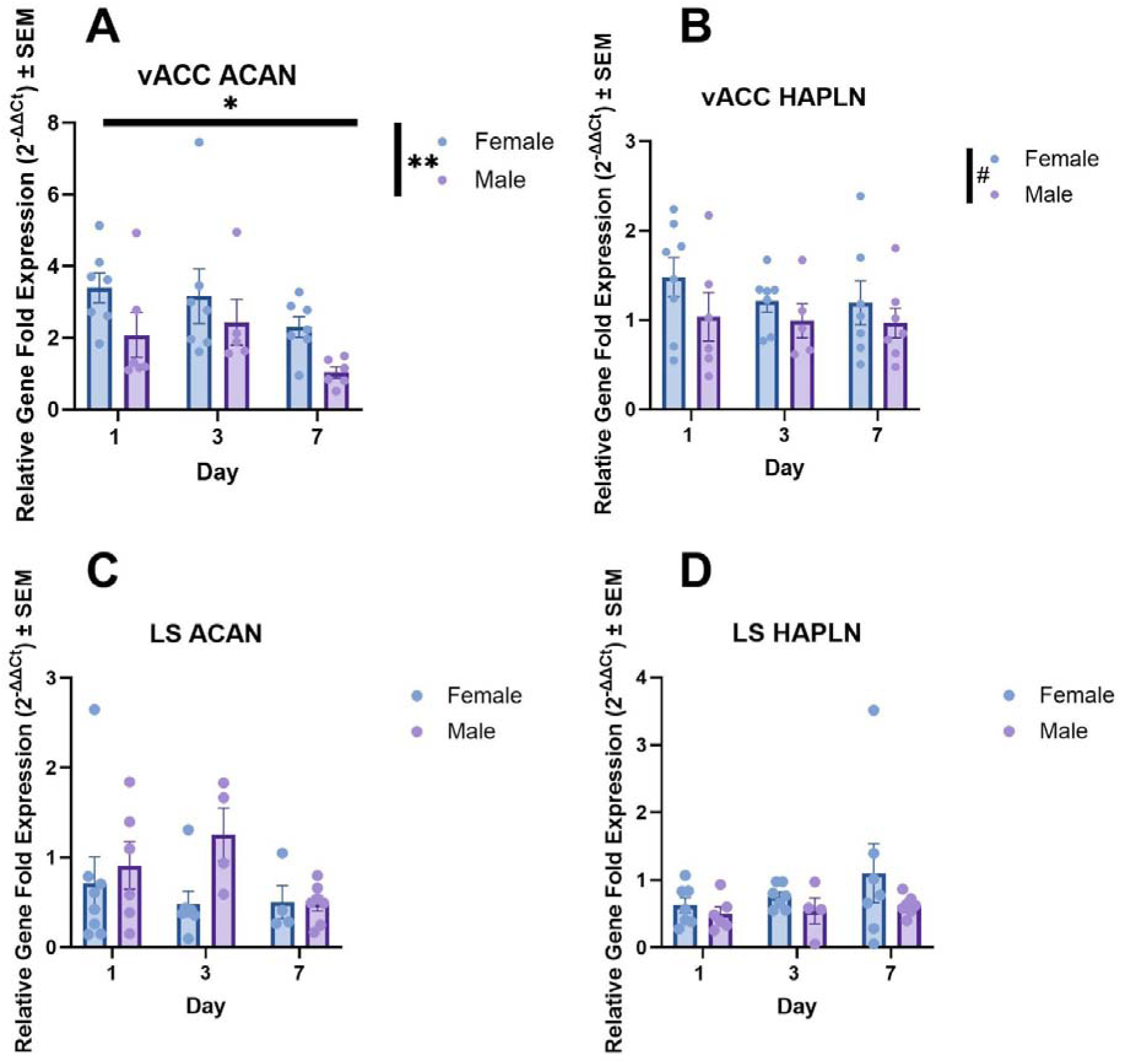
Clear sex differences in structural plasticity are present in the ventral anterior cingulate cortex (vACC), but not the lateral septum (LS), alongside a significant decrease in PNNs across pair bond establishment. A) ACAN expression decreased over the course of pair bonding in the vACC (p=0.030), but there was a noted sex difference with females displaying higher relative levels of vACC-ACAN mRNA across the first week of bonding (p=0.004). B) In contrast, HAPLN mRNA expression displayed a nonsignificant trend of higher expression in females, although there was still no significant difference over 1-wk of bonding (p=0.086). C) Finally, ACAN and HAPLN expression were not found to differ significantly as a function of either day and sex in the LS. **#= p<0.1, *= p<0.05, ** = p<0.01, *** = p<0.001**

#### vACC-OXTR and vACC-AVPR

There was a significant main effect of sex on OXTR mRNA expression in the vACC F (1, 32) = 8.165, p=0.007, Fig. 3A). Females, on average, expressed OXTR mRNA at higher levels than males. However, neither the effect of pair bonding day, nor the interaction between day and sex on relative OXTR mRNA expression were significant (see Supplemental Table 1). We detected a significant sex effect in which males displayed higher levels of AVPR mRNA in the vACC than females (F (1, 33) = 8.910, p = 0.005, partial η2 = 0.21 Fig. 3B), but neither the interaction between sex and day, nor the main effect of day itself were significant (see Supplemental Table 1). Females had significantly higher ratios of OXTR/AVPR than males (F (1, 31) = 7.816, p= 0.009, partial η2 = 0.20 Fig. 2C). No additional main effects or interactions were significant. Ultimately, sex differences were seen in all measures of receptor plasticity, but no effect of pair establishment day was apparent across any measure of receptor mRNA (see Supplemental Table 1 for statistics).

**Figure 3.**
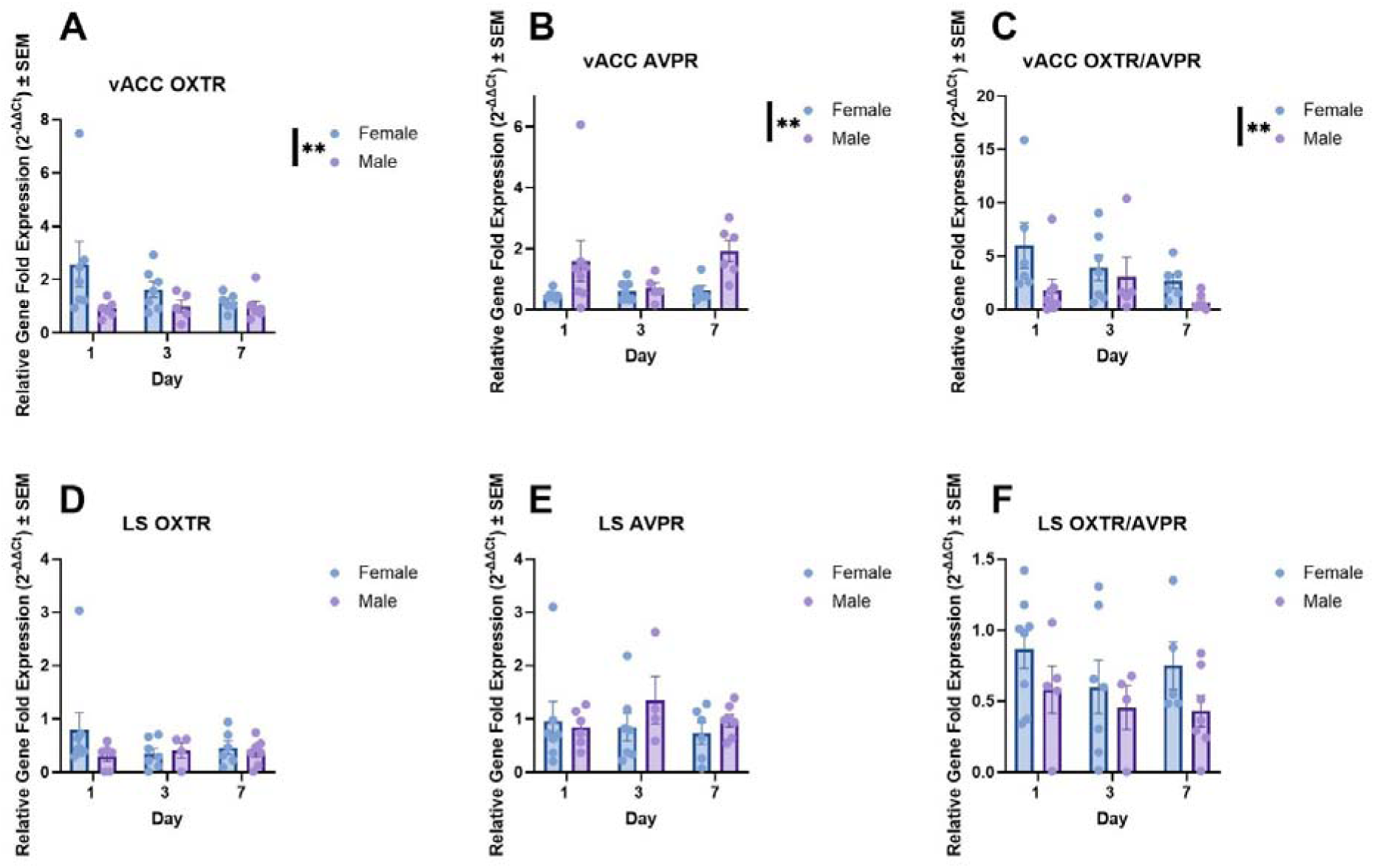
Ventral Anterior Cingulate Cortex (vACC), but not Lateral Septal tissue (LS), displayed differences in OXTR to AVPR mRNA between sexes, but no other significant findings were detected. A) A significant sex difference in which females displayed higher levels of relative OXTR gene expression in the vACC was detected (p=0.007), alongside B) another sex difference in which males displayed higher levels of AVPR in the vACC (p=0.005). C) This sex difference was also reflected in the ratio between OXTR to AVPR in the vACC where females, on average, displayed higher ratios in the vACC (p=0.009). D-F) Conversely, no significant sex differences were detected in OXTR, AVPR, or the ratio of the two receptor types expressed in the LS. **\*= p<0.05, ** = p<0.01, *** = p<0.001**

#### LS-OXTR, AVPR, HAPLN, and ACAN

All data (with the exception of the LS-OXTR/AVPR ratio) were transformed due to normality and heteroscedasticity violations. No significant effects of day or sex were detected across all gene expression investigations in the LS (see Supplemental Table 1 for statistics).

#### Gene Expression and Associations with Affiliative and Aggressive Behavior in Early and Late Bonding

Males with neural tissue extracted 24 hours post-introduction displayed a significant positive correlation between vACC AVPR mRNA expression and introductory chasing behavior (R= .977, p<.0001, Fig. 4A). Females with neural tissue extracted 24 hours post-introduction also displayed a significant positive relationship between the vACC OXTR/AVPR and introductory chasing behavior (R= .979, adj. p<.0001). Nesting distance was negatively associated with vACC ACAN mRNA expression, meaning smaller nesting distances tended to exist alongside higher levels of ACAN expression (R= .916, adj. p=0.001). Statistically significant associations between mRNA measures persisted in females after p-value corrections. Namely, LS OXTR was positively associated with LS AVPR and LS ACAN expression (R= .937, adj. p= .001 ; R= .980, adj. p<.0001).

**Figure 4.**
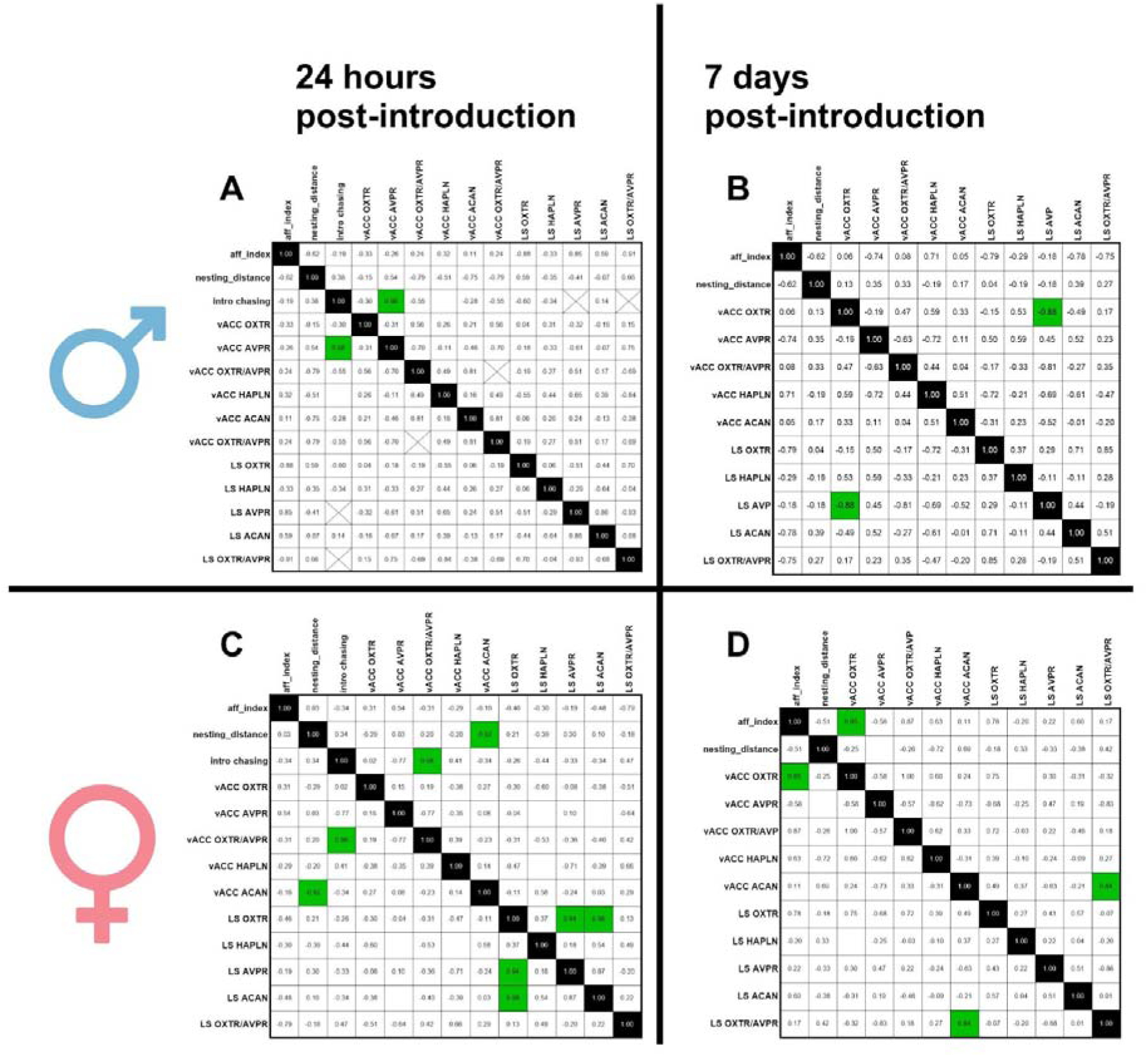
Correlation matrices of behavior and relative mRNA expression in the ventral anterior cingulate (vACC) and the lateral septum (LS) between males and females in 24 hours post-introduction (Group 1) and 7 days post-introduction (Group 3). Early AVPR associations with aggressive behavior shift to female-driven correlations between vACC OXTR measures and affiliation by late bonding. **A)** Males who were sacrificed 24 hours post-introduction to a female pair mate displayed a significant positive relationship with vACC AVPR mRNA expression with introductory chasing behavior. **B)** In males that were sacrificed 7 days post-introduction to a female pair mate displayed no significant relationships between behavior and any mRNA measures. **C)** Meanwhile, females who were sacrificed 24 post-introduction to a male pair mate displayed a contrasting but significant positive association between vACC OXTR/AVPR ratios and aggressive chasing behavior. Furthermore, a significant negative association between vACC ACAN mRNA expression and nesting distance in females, in which smaller nesting distance represents greater affiliation at 24 hours post-introduction, was detected. **D)** In comparison, females sacrificed 7 days post-introduction appeared to drive significant positive correlations between vACC OXTR mRNA and the index of affiliative behavior. **Green cells within the matrices indicate significant correlations after Benjamini-Hochberg correction.** Created with Biorender.com.

Males with neural tissue extracted 7 days post-introduction displayed a significant negative relationship between vACC OXTR mRNA expression and LS AVPR mRNA expression (R= -.88, adj. p<.01). Females with neural tissue extracted 7 days post-introduction exhibited one positive significant relationship between the affiliative index and mRNA measures. Namely, the affiliative index was associated with vACC OXTR (R=.847, adj. p= .008). Additionally, LS OXTR was positively associated with vACC ACAN mRNA expression (R=.996, adj. p<.0001). No additional significant findings were detected after p-value corrections.

### 1.3.3 Immunohistochemistry of Perineuronal Nets (PNN)

As a comparison to the qPCR, we also used immunocytochemistry to examine PNN+ neurons in the vACC, ventral LS and dorsal LS In a subset paired (n=6) and unpaired (n=6) animals, we compared the immunohistochemistry cell counts of PNNs in three brain areas: vACC, dLS, vLS. Across all comparisons, Welch’s t-tests were used to compare unpaired and paired individuals. Individuals in each group were sex balanced. To evaluate correlative comparisons, a Pearson’s correlation coefficient was used to evaluate the strength and direction of associations. No comparisons of PNN+/PV+, PNN+ only, PV+ only cells were significantly different between unpaired and paired animals (see Figure 5).

**Figure 5.**
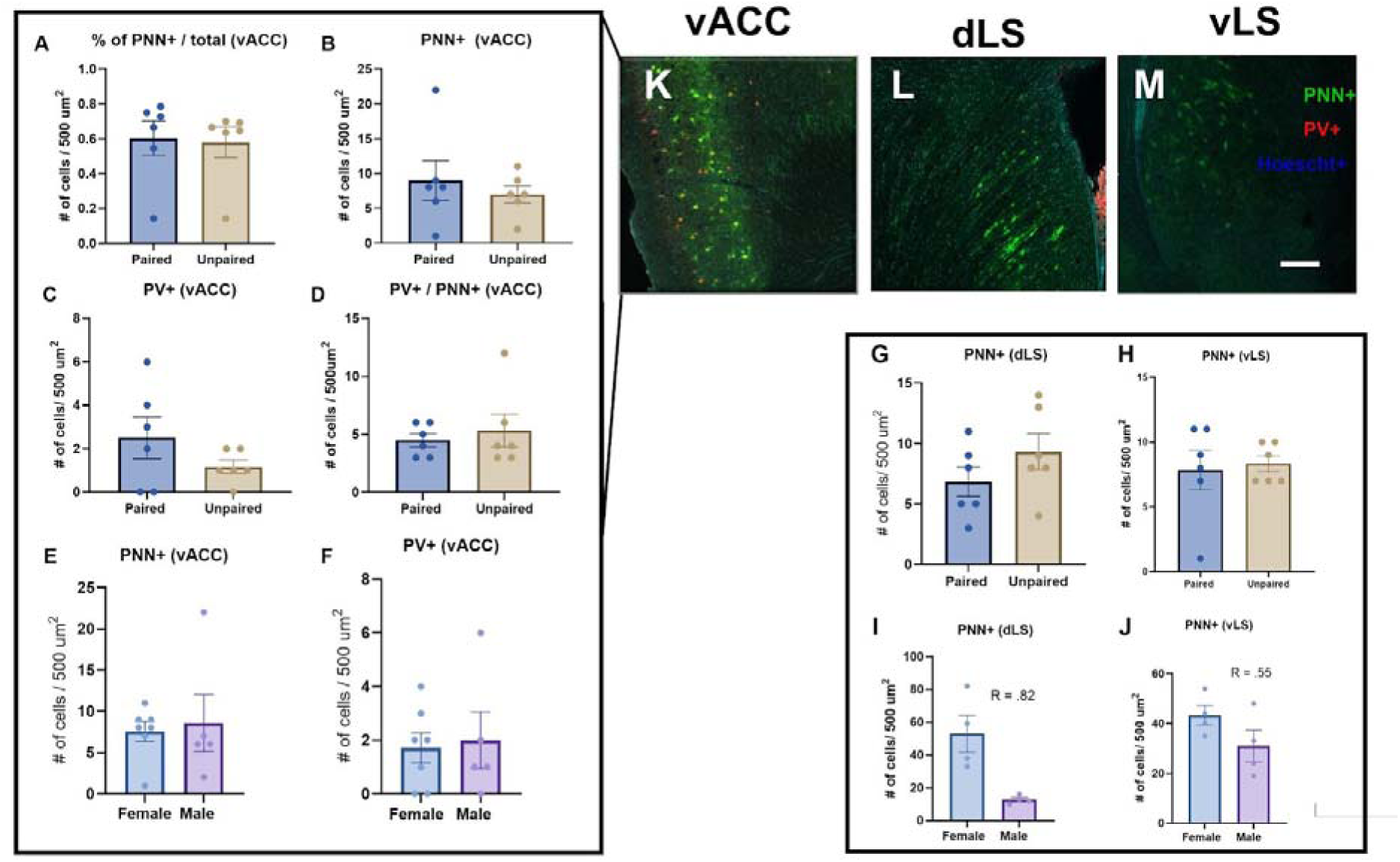
IHC analysis of PNN+ and PV+ cells did not detect differences between unpaired and paired individuals but inconsistent with qPCR, a sex difference was found in the LS. **A-H)** No significant differences in PNN numbers in the vACC or LS. **A-D)** No significant differences were apparent between unpaired individuals in the vACC. Neither a difference in the proportion of PNN+ cells to total cells **(A)**, raw PNN+ numbers **(B)**, raw PV+ numbers **(C)**, nor a ratio of these cells **(D)** were noted. **E-F)** No significant sex differences in PNN+ **(E)** or PV+ **(F)** cell numbers were noted in the vACC. **G-H)** In the LS, no significant differences in PNN+ cell numbers were detected. **I-J)** Notable sex differences, with medium to large effect sizes, were noted in each subdivision of the LS: dLS (p<0.001) and vLS (p<0.001), with females displaying higher numbers of PNN+ cell numbers. **K-M)** Representative images of IHC in the **(K)** ventral anterior cingulate cortex (vACC), **(L)** dorsal lateral septum (dLS), and **(M)** the ventral lateral septum (vLS). Each representative image displays three stains: PNN+ cells (green), PV+ cells (red), and a nuclear stain (blue). *Note that the vACC was the only area to display PV+ staining, as such comparisons for paired and unpaired animals and males and females include PV+ cell counts for this area only. **Scale bar = 200um.**

Females exhibited higher numbers, sometimes 3-fold increases, in the number of PNN+ positive cells in the dLS (t(11.49) = 6.46, p <0.001, partial η2 = 0.78; (Fig. 5G)). A similar sex difference with females possessing higher numbers of PNN+ cells in the vLS was also present (t(11,79) = 8.16, p <0.001, partial η2 = 0.85; Fig. 5H). Beyond this, no other notable or significant comparisons were detected. Differences in IHC and qPCR results are discussed below.

## 1.4 Discussion

Social species undergo clear socio-behavioral changes, such as pair bonding, that serve to increase fitness and reproduction. However, the complex underlying biological changes that are associated with these behavioral shifts have not been fully studied. In this study, we aimed to both characterize the timing of behavioral changes that California mice undergo as they establish a monogamous bond and examine structural and receptor plasticity in both sexes across a seven day period. We detected a statistically significant reduction in PNN-related gene expression (ACAN) of both sexes combined across the first week of pair establishment in vACC. PNNs have repeatedly been implicated in the fine tuning and modulation of social behavior across species. Most notably, across gestation, pup rearing, and weaning, PNNs dramatically change in the medial preoptic area of female mice across offspring rearing (Uriarte et al., 2020), with notable reductions in PNN coinciding with behavioral and hormonal shifts (e.g. parturition and cessation of lactation). Furthermore, as previously mentioned, song learning in social bird species (zebra finches) appears to coincide with higher PNN levels present after seasonal song learning periods when song crystallization has occurred and song variation has decreased (for review see Balthazart, 2023). In this investigation, we discovered an associative link between PNN production and the onset of monogamous behavior. In addition, we noted sex specific differences in structural and receptor plasticity in the vACC, a key brain area in the control of social behavior. In order to detect this difference, we utilized a two-pronged approach which included both qPCR and IHC measurements of two key brain areas–the vACC and LS. Both methods have been used in tandem to evaluate the dynamic remodeling and general presence and localization of PNNs, respectively (Kim et al., 2017). Our analysis heavily emphasizes the qPCR data, as the sample size for our IHC cohort was smaller (n=12).

### 1.4.1 Rapid Shifts in Aggression Followed by Gradual Changes in Affiliation

Using daily homecage observations, we were able to evaluate the nonspecific aggressive and partner-specific affiliation changes that occur across the pair bonding establishment period in greater detail than previous investigations. We discovered an expeditious decline in aggressive tendencies from Day 0 (Introduction) to Day 1. Succeeding this, we documented a graded increase in affiliative tendencies (huddling and social grooming), which peaked at Day 3 of pair establishment. Furthermore, other affiliative behaviors, namely nesting separation, declined across the entire week-long establishment period. Alongside our behavioral investigations, we also recorded USVs during homecage observations and noted a significant decrease in both sustained vocalizations and sweeps across the pair bond establishment period. These decreases in vocalization number coincided with olfactory investigation decreases seen from Day 0 to Day 1. These findings further solidify that the California mouse’s selective, long-lasting bond takes longer to form than other species with similar mating structures (prairie voles, see Introduction for more information), and may be a better suited model for investigating plasticity fluctuations associated with long term behavioral alterations.

Alongside these clear aggressive and affiliative behavioral shifts, we also investigated the relationships between all behaviors measured and their associations with vocalization features. We discovered a clear positive association with olfactory investigation duration and vocalization counts. This association is similar to that found in previous investigations of California mice in which sweep number is positively correlated with sniffs (Pultorak et al., 2018). For more information on behavioral and vocalization correlations see Supplemental Figure 1.

### 1.4.2 Structural Plasticity Increases Across Pair Bonding

ACAN is an essential component of PNNs and is required for PNN function (Suttkus et al., 2014). We found a significant reduction in relative ACAN mRNA expression from Day 1 to Day 7 of the pair bond establishment period in both sexes in the vACC. No changes were detected in HAPLN possibly because HAPLN appears to be less responsive to environmental changes as it fulfills multiple roles, whereas ACAN is vital in producing the dynamic changes seen in PNN (Rowlands et al., 2018; Kwok et al., 2010). The decrease in ACAN indicates a potential role of PNNs in the production of the behavioral changes associated with formation of monogamous bonds and, potentially, preparation for offspring rearing. Because this change was detected in the vACC and not the LS using qPCR measurements, we suspect that PNN change in this area may represent changes to individual social salience networks related to a relatively long term change in the social environment. We speculate that upon cohabitation, pair mates are learning how to identify their mate such as via vocalization features (Pultorak et al., 2017;Warren et al., 2025), coordinate behaviors for optimized pair living, and begin establishing neural circuits and behaviors conducive to cooperation in raising young. PNN mRNA decreases may signal an increase in the flexibility of these circuits that allow for such cue learning. This idea is supported by the vACC’s role in social decision-making processes across species. In rodents and nonhuman primates, the ACC plays a key regulatory role in social cue salience and valuation. In particular, the ACC is involved in accurate valuation judgements (Rudebeck et al., 2006) and social cue sensitivity (Swain et al., 2023). Furthermore, a similar role of the vACC has been established in humans in which the vACC regulates salience and value judgements (e.g. Rigney et al., 2018). With this in mind, we speculate that the PNN-proxy mRNA reduction detected in the vACC may function to regulate the learning of new social cues. This could allow for more flexible learning as pair mates begin to form a unique, cooperative dyad (Rieger et al., 2018, Malone et al., 2023). It is possible that this transient decrease in PNN-proxy mRNA could be followed by a learning solidification process in which PNN mRNA expression increases. This would likely occur many days to weeks after the initial pair bond is fully established, as seen in previous investigations of learning/social state changes (Cornez et al., 2020; Uriarte et al., 2020). This is further supported by the significant association between nesting distance and vACC ACAN mRNA expression in females paired for 24 hours. Other studies with the California mouse also reveal changes in vocalizations over the first week of bonding such as a lengthening of the SVs (sustained vocalizations) (Pultorak et al., 2017), suggesting that there is a suite of traits that take longer to adjust after initial pair introduction. To test PNN’s role on pair bonding behavior, future investigations could use chABC intracerebral injections, which digest PNN components, into the vACC to evaluate the role of these structures in behavioral plasticity across this timeframe (Banjeree et al., 2017).

### 1.4.3 Sex Differences in mRNA expression in the vACC and LS

We detected numerous sex differences in mRNA expression within the vACC, a cortical region, but not the expected difference in the LS, a subcortical region. Significant sex differences were detected in every gene expression measure in the vACC: OXTR, AVPR, ACAN and a nonsignificant trend in HAPLN. Females possessed higher levels of OXTR, ACAN, and a nonsignificant trend in HAPLN, but lower levels of AVPR mRNA. Sex differences in cell connectivity and function have been noted in the ACC, specifically with males exhibiting heightened dendritic arborization and increased cue-associated activation in attention tasks (Markham et al., 2002; Liu et al., 2012), but to our knowledge, an investigation into sex differences in receptor and structural plasticity has not occurred. Previously, OXTR binding density was assessed in the California mouse in the broad cingulate cortex, but binding affinity for OXTR was higher in males (Insel et al., 1991). Our results do not align with this finding, possibly because we exclusively focused on the vACC rather than including the dorsal anterior cingulate cortex, which is implicated in a suite of different behaviors.

The sex differences we discovered in vACC-OXTR and vACC-AVPR mRNA expression align with classical investigations of pair bonding and other behaviors in male and female prairie voles. In such studies, both AVP and AVPR promote facilitation of pair bonding in males, while OXT and OXTR signalling/modulation tend to increase pair bonding behaviors in females (Cho et al., 1999; Young et al., 2001; Lim et al., 2004). These tendencies have also extended into the human literature on strong social attachments (Walum et al., 2008; 2012), but this relationship in humans is more complex. Our study further supports the sex-specific roles of AVPR and OXTR in social attachment in mammalian species. OXT and AVP also support other behaviors advantageous to biparental species. For example, AVP immunoreactivity has been positively associated with paternal behavior in California mice (Bester-Meredith & Marler, 1999) and generally positive associations with paternal behavior in other species (Guoynes & Marler, 2020), but results vary, as in the case of nest-building in *Peromyscus* (Bendesky et al., 2017).

We also discovered significant correlations with medium to large effect sizes between behavior and neuropeptide receptor mRNA at different points across pair establishment. Most notably, vACC-AVPR mRNA in males and its ratio with vACC-OXTR in females correlate positively with chasing behavior at pair introduction (Supplemental Figure 3). We speculate that AVPR in the vACC might influence the social salience of aggressive encounters in this highly territorial species (Rigney et al., 2018; Rigney et al., 2022). In addition, females displayed correlations with vACC-OXTR mRNA and the affiliative index calculated for Day 7, further supporting the importance of OXT signalling in the formation of female bonding (Supplemental Figure 3). Males did not display this vACC-OXTR mRNA correlations. As such, the vACC should be considered a relevant node in the future investigation of social behaviors such as pair bonding.

While sex differences in OXTR and AVPR utilization are well-supported by past literature, sex differences in the cortical expression of PNNs are underexplored. Sex differences in PNN have been noted in a few studies in areas with highly sexually dimorphic function/anatomy, such as nuclei of the hypothalamus in mammals (Ciccarelli et al., 2021; Zhang et al., 2021), and in cortical song learning nuclei in zebra finches (Cornez et al., 2015). The current study further supports cortical sex differences in PNNs, as we discovered that females expressed significantly higher levels of ACAN mRNA (and a nonsignificant trend of higher HAPLN mRNA) in the vACC. It is unclear why males express lower levels of vACC-ACAN mRNA. There are some behavioral differences that would naturally occur at this time point. Males experience a greater behavioral shift from post-weaning territorial, solitary living to cooperative pair living (Zhao et al., 2020; Ribble, 1992). Males disperse and establish territories near their natal territory first, and later females disperse farther to find a mate (Ribble et al., 1992). Males display slightly higher levels of contact-based aggression, differences in territorial scent-marking patterns, and earlier aggressive signals in male-male encounters, though both sexes display territoriality (Kuske et al., 2024; Bester-Meredith et al., 2005; Becker et al., 2018; 2010; Rieger et al., 2018). As such, it is possible that males require heightened behavioral and neural flexibility after weaning in order to balance mate acquisition and territory defense. Meanwhile, females, which are territorially aggressive, but to a lesser degree, and already exhibit higher levels of affiliation in same-sex interactions prior to pair-bond formation may not require as much flexibility (Kuske et al., 2024; Bester-Meredith et al., 2005; Hammond et al., 2025). These behavioral changes, alongside preparation for dramatic behavioral and physiological fluctuations associated with future pregnancy and offspring rearing could account for sex differences seen in the vACC.

We did not detect significant changes in OXTR or AVPR mRNA in the vACC or LS across the pair establishment period, which did not align with our original hypothesis that predicted positive and/or negative changes in receptor mRNA. It is possible that using unmatched pairs (pairs that are dramatically different in their approach/avoidance tendencies), could reduce sample variability in future studies, which may allow for significant changes in receptor plasticity to be discovered (Monari et al., 2021). Alongside this, PNNs that form extracellular matrices around neurons regulate receptor plasticity (Sorg et al., 2016). The presence of PNNs on cells inhibits receptor movement and integration within the cell membrane. As such, changes in OXTR and AVPR mRNA expression could be preceded by reductions in PNNs. After which, changes in receptor densities could be more easily enacted on the cell membrane. This study may not have captured the extended timeframe needed for receptor changes after pair introduction.

In the LS, we did not encounter the same sex differences documented in OXTR and AVPR expressing cells of the vACC. However, we speculate that this occurred because of sampling the entire LS, rather than separating the ventral, intermediate, and dorsal subregions, which possess sequestered subpopulations of cells that differ between sexes in number and cell type (Olivieria et al., 2019; 2021; Tsukahara et al., 2004). In an older study of the California mouse, males possessed higher OXTR binding densities in the dLS (Insel et al., 1991), but we did not detect this in our analyses of the full LS. This suggests this sex difference may be limited to the dLS. Future studies could attempt microdissection of these subregions, if possible, to investigate neuropeptide receptor gene expression and sex differences in each distinct and important area.

### 1.4.4 Associations in the PNN system of the vACC and LS

Another correlation of note is that between vACC-ACAN and nesting distance in females sacrificed 24 hours after pair introduction. This correlation was unexpectedly negative, meaning higher ACAN expression was associated with shorter nesting distances at 24 hours post-introduction. We can only speculate, but it raises the possibility that an increase in ACAN expression as a result of co-nesting could be used to stabilize new social learning structures in the vACC. Co-nesting may be especially important, as it may signal a later stage solidification of the California mouse’s lifetime bond and is possibly reflected in ACAN’s decrease across the first week of pair bond establishment (Stoppel et al., 2024; Khadraoui et al., 2022; Supplemental Figure 2). The function of the association between the timing of PNN densities and changes in behavior are unclear.

### 1.4.5 Discrepancies in IHC and qPCR Outcomes

Our differential results between IHC and qPCR highlight the technique-specific benefits and pitfalls that bolster support for the use of both methodologies in tandem to secure a more refined picture of the dynamic PNN remodeling that occurs in region-specific patterns. Using qPCR, we may have had greater sensitivity to detect sex differences in vACC PNN measures across a 1 week timeframe, but IHC revealed sex differences in PNN+ cell numbers across LS subregions (vLS and dLS) that qPCR failed to detect. qPCR allowed us to detect fluctuations in PNN within large homogenates of neural tissue, which supplies us with greater understanding of the dynamic nature of this system that occurs across time (for reviews see Lorenzo Bozzelli et al., 2018; Carulli et al., 2021; Santos-Silva et al., 2024). In contrast, given the localization specificity of IHC, we were able to detect subregion-specific sex differences in PNN numbers that are not detectable using qPCR. It is not surprising that few differences were found using IHC (due to simply counting cell numbers and small sample size) but the sex differences found in the LS are likely due to the sexual dimorphic behavioral utilization of the LS between the sexes. Nonetheless, future studies could gain more from such IHC analyses by using a more qualitative approach looking at the different PNN qualities (e.g. number of processes covered, relative fluorescence, etc.). However, both techniques revealed sex differences in which females appeared to possess or express higher cell numbers or mRNA quantities of PNN-related measures. This propels us to suggest the dual usage of these techniques (as seen in previous studies) to investigate this complex system (Kim et al., 2017).

### 1.4.6 Conclusions

Our study provides novel associative evidence that pair bonding behavior, including co-nesting, could be regulated by structural plasticity mechanisms, such as PNNs. Further manipulations of PNNs (e.g. intra-cerebral chABC injection to the vACC), could provide support for this mechanism’s causal role in social behavior beyond maternal care and song crystallization. Our study also further expands our understanding of sex-specific relative expression of nonapeptide receptors in a cortical area (vACC), which has previously been underexplored. Future studies should include this area in further investigations of the role of structural and receptor plasticity changes on social behavior, including monogamous pair bonding.

### 1.4.7 Perspectives and Significance

This study provides the first evidence that changes in structural plasticity, in the form of PNNs, parallel the development of long lasting monogamous bonds in both sexes. Furthermore, this study provides novel evidence supporting highly differential OXTR and AVPR expression in specifically cortical tissue between males and females. This is a previously unknown sex difference that should facilitate further study in the differences of receptor expression between the sexes in cortical areas. The vACC in particular should now be considered potentially important for bonding, as this study reinforces its involvement. Furthermore, future investigations in sex differences should focus on the use of structural plasticity (PNNs) to modulate social salience, learning, and decision-making.

## Acknowledgments

A special thanks to Kelci Cox for her additional contributions to the training and use of DeepSqueak on *Peromyscus californicus* vocalizations. We would like to thank Changjiu Zhao and Lauren Riters for their expertise and qPCR equipment. We would also like to thank our excellent animal care staff at UW-Madison. This study was supported by the following grants: UW-Madison Schwartz Fellowship, Animal Behavior Society MSN278502, and NSF 1946613.

## Declarations

### Ethics approval

Procedures used in this study were approved by the University of Wisconsin– Madison College of Letters and Sciences Institutional Animal Care and Use Committee (Protocol L005447).

### Consent for publication

not applicable.

### Availability of data and materials

Data is available upon request.

### Funding

This study was supported by the following grants: UW-Madison Schwartz Fellowship, Animal Behavior Society MSN278502, and NSF 1946613.

### Competing Interests

The authors declare that they have no competing interests.

### Clinical Trial Number

not applicable

### Consent to publish declaration

not applicable

### Ethics and Consent to Participate declarations

not applicable

## Supplemental Information

**Supplemental Figure 1.**
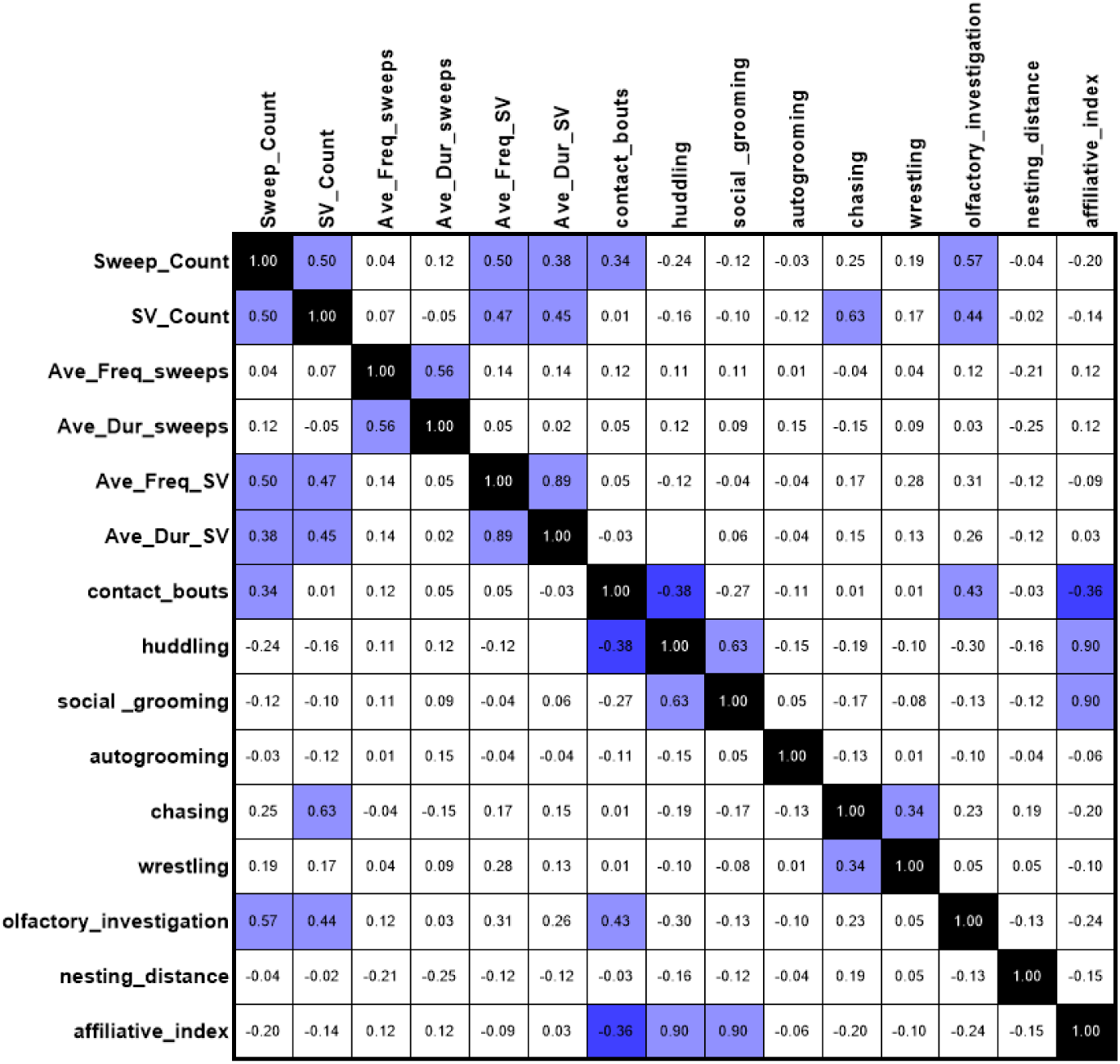
Correlation matrix between behavior variables and vocalization features. Blue shading of cells indicates a persistent significant correlation after a Benjamini Hochberg multiple comparison p-value correction.

**Supplemental Table 1.** Nonsignificant RT-qPCR Two-Way ANOVA Results. F-statistics for each two-way ANOVA and its interaction is listed for each mRNA measure in each brain area (vACC and LS). * indicates statistical significance.

|  | Effect of Day | Effect of Sex | Interaction |
| --- | --- | --- | --- |
| vACC ACAN | F (2, 32) = 3.926* | F (1, 32) = 9.799* | F (2, 32) = 0.5395 |
| vACC HAPLN | F (2, 34) = 0.3022 | F (1, 34) = 3.130 | F (2, 34) = 0.2152 |
| vACC OXTR | F (2, 32) = 1.195 | F (1, 32) = 8.165* | F (2, 32) = 1.692 |
| vACC AVPR | F (2, 33) = 1.851 | F (1, 33) = 8.910* | F (2, 33) = 1.873 |
| vACC OXTR/AVPR | F (2, 31) = 1.821 | F (1, 31) = 7.816* | F (2, 31) = 0.6704 |
| LS ACAN | F (2, 30) = 1.369 | F (1, 30) = 2.680 | F (2, 30) = 1.371 |
| LS HAPLN | F (2, 32) = 0.7669 | F (1, 32) = 2.254 | F (2, 32) = 0.1027 |
| LS OXTR | F (2, 32) = 0.2520 | F (1, 32) = 1.606 | F (2, 32) = 1.198 |
| LS AVPR | F (2, 31) = 0.3691 | F (1, 31) = 1.859 | F (2, 31) = 0.5282 |
| LS OXTR/AVPR | F (2, 30) = 0.7943 | F (1, 30) = 3.620 | F (2, 30) = 0.1533 |

### Primer design

Primer pairs were designed via aligning mus musculus genome to the peromyscus genus using NCBI Nucleotide BLAST. Aligned sequences were then transferred into NCBI primer BLAST to create numerous primer pairs for the Peromyscus aligned sequence with a desired amplicon of 200bp or smaller. All primers were designed to run at a melt temperature of 58°C. Primer pairs were validated via single peak amplification and standard curve efficiency analysis using cDNA synthesized from extracted RNA from neural tissue samples of interest (vACC and LS). All cDNA was diluted to the lowest cDNA concentration (evaluated via NanoDrop 2000 (ThermoScientific)) prior to validation and experimental runs.

**Supplemental Figure 2.**
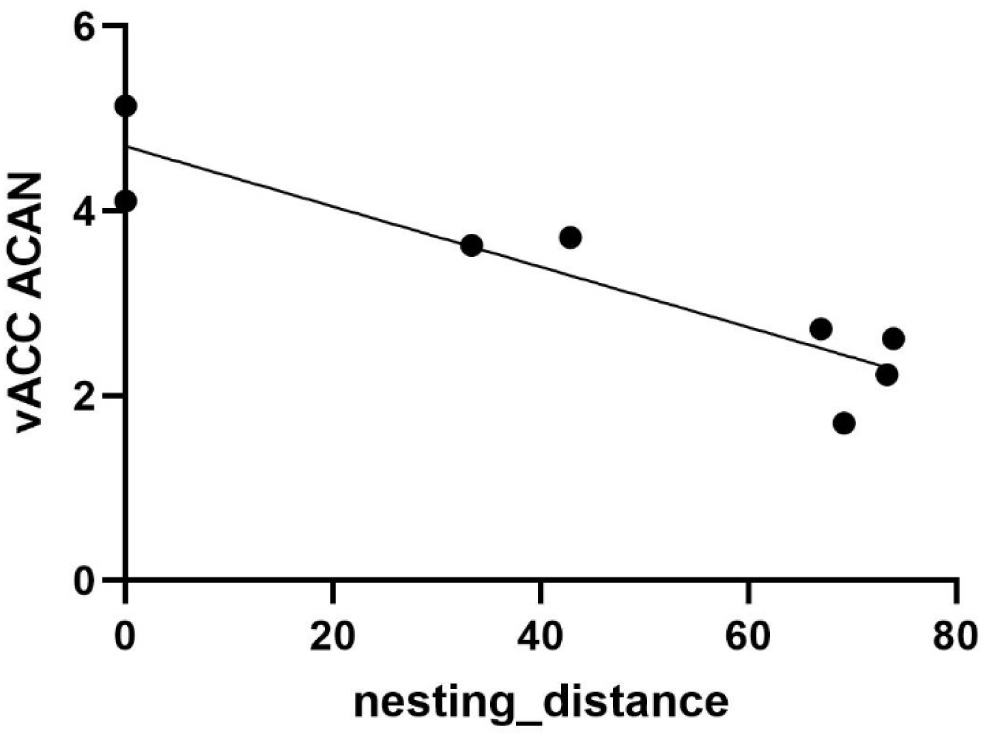
vACC ACAN is negatively correlated with nesting distance in females sacrificed 24 hours after pair introduction.

**Supplemental Figure 3.**
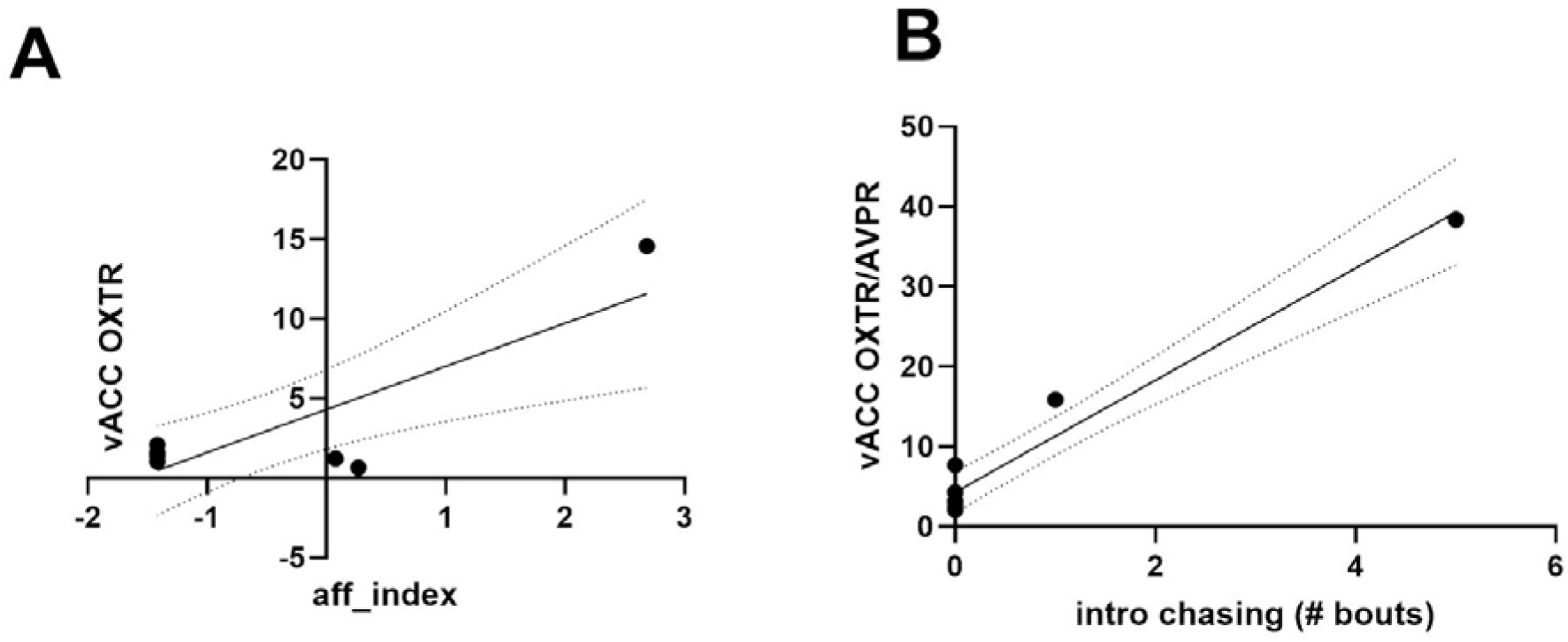
Behavioral and qPCR correlations for females 24 hours after introduction and 1 week after introduction. **A)** Linear correlation with 95% CI of the positive relationship between vACC OXTR mRNA expression and the behavioral affiliative index on Day 7 of pair establishment (R = 0.85). **B)** Pearson correlation line with 95% CI of the positive relationship between introductory chasing bouts and vACC OXTR/AVPR mRNA expression 24 hours after pair introduction (R = 0.98).

**Effect Size Considerations (partial η2) :**

Small: 0.01-0.05

Medium: 0.06-0.13

Large: 0.14+

**Effect Size Considerations (R-squared)**

Small: 0.02-0.12

Medium: 0.13-0.25

Large: 0.26+

## Notes

### Competing Interest Statement

The authors have declared no competing interest.

